# The systemic activin response to pancreatic cancer: Implications for effective cancer cachexia therapy

**DOI:** 10.1101/343871

**Authors:** Xiaoling Zhong, Marianne Pons, Christophe Poirier, Yanlin Jiang, Jianguo Liu, George E. Sandusky, Yunlong Liu, Guanglong Jiang, Marion Couch, Leonidas G. Koniaris, Teresa A. Zimmers

## Abstract

Pancreatic ductal adenocarcinoma (PDAC) is a particularly lethal malignancy with high rates of cachexia. Serum activin correlates with PDAC cachexia and mortality, while activin administration causes cachexia in mice. We studied activin in human tumors and in mice with orthotopic or genetic PDAC. Cachexia severity correlated with activin expression in tumor lines. Activins were expressed in both cancer and tumor stromal cells, but also in organs in murine PDAC cachexia. Tumor cells expressed activin-βA, or *Inhba*, while organs expressed both activin-βA and activin-βB, or *Inhbb*. PDAC elicits activin expression; PDAC conditioned medium induced activin and atrophy of myotubes. Treatment with the activin trap, ACVR2B/Fc, reduced cachexia and prolonged survival in mice with activin-low tumors, and reduced cachexia in activin-high tumors, without affecting activin expression in organs. Mice expressing dominant negative ACVR2B in muscle were protected for weight loss but not survival. Overall our results indicate that PDAC induces a systemic activin response, leading to cachexia, and that activin targets might include organs beyond muscle. Targeting of both tumor-derived and host-derived activins could improve cachexia therapy.

## Introduction

Cachexia is a multifactorial syndrome characterized by an ongoing loss of skeletal muscle mass (with or without loss of fat mass) and other clinical manifestations of cachexia include anorexia and fatigue (Blum et al, 2014; Fearon et al, 2011). The fact that simply increasing of food intake or nutritional support is inadequate to reverse the cachectic state indicates a primary pathology in wasting organs. This debilitating state negatively affects the ability of patients to tolerate surgical resection of tumor and anticancer therapy such as chemotherapy and radiotherapy. Indeed, cachexia has become an obstacle for the successful treatment of cancer and may significantly contribute to cancer death. However, there is no established effective therapeutics to treat this disease.

Activins are members of the transforming growth factor-beta (TGF-β) superfamily of signaling proteins that share structural similarities but diversify in biological activities or functions. The principal function of inhibin is regulating gonadotrophin, whereas activin plays multiple biological roles including the regulation of development, homeostasis, tissue repair, inflammation, fibrosis, and tumorigenesis and progression, in addition to the FSH-stimulating role (Asashima et al, 2000; Hedger & Winnall, 2012; Kim et al, 2000; Loomans & Andl, 2014; Werner & Alzheimer, 2006).

Activins are composed of two polypeptide subunits and each subunit is encoded by a separate gene: subunit βA (gene *Inhba*), βB (*Inhbb*), βC (*Inhbc*), and βE (*Inhbe*). The two subunits are linked by a single disulfide bond. Homodimer of two βA or two βB gives rise to activin A (βAβA) and activin B (βBβB), respectively; heterodimer of one βA and one βB was named activin AB (βAβB) (Ling et al, 1986a) (Ling et al, 1986b). βC (G et al, 1995) and βE (Hashimoto et al, 2002) were discovered more recently and their biological roles are largely unknown.

Activin A is the best characterized member of the activin subfamily involved in many biological and pathological processes as stated earlier. Deregulated activin A expression is often observed in various malignancies (Kleeff et al, 1998; Loomans & Andl, 2014; Wildi et al, 2001). Overexpression of activin A in the tumor tissues (Kleeff et al, 1998) or elevated blood activin A levels were found in patients with pancreatic cancer (Togashi et al, 2015) and other cancer types (Harada et al, 1996; Hoda et al, 2016; Leto et al, 2006; Loumaye et al, 2015; Terpos et al, 2012), suggesting a potential “endocrine” effect of activin A on the host organs.

The high prevalence (~ 85%) of cachexia has significantly contributed to PDAC death. Despite this recognition, the mechanism underlying the PDAC cachexia is still incompletely understood. Several studies correlated the elevated plasma activin A level with cachexia induced by PDAC and other cancer types and with shorter survival (Chen et al, 2016; Chen et al, 2014; Loumaye et al, 2015; Loumaye et al, 2017; Togashi et al, 2015). In preclinical models, a direct causal link between activin A and muscle wasting/cachexia demonstrated that elevating circulating activin A by injecting the activin A gene-carrying virus vector into the leg muscle of tumor-free mice reduced body weight (Chen et al, 2016; Chen et al, 2014).

Our group and others have shown that inhibiting myostatin and related ligands, such as activin and GDF11, by systemic administration of the activin receptor extracellular domain/Fc fusion protein, ACVR2B/Fc potently inhibited muscle wasting in both colon-26 and Lewis lung carcinoma cachexia mouse models (Benny Klimek et al, 2010; Zhou et al, 2010). These studies established that targeting myostatin-family ligands using ACVR2B/Fc or related molecules is an important and potent therapeutic avenue in cancer cachexia.

Given the deregulation of activin A seen in many cancer types including PDAC and the potentially systemic impact of elevated blood activin A on the host organ biology, in the present study, we aimed to investigate whether activin mediates cachexia in murine models of PDAC cachexia as well as in patients and to evaluate the therapeutic utility of blocking the ACVR2B-mediated signaling pathway. Our study may lead to identification of novel candidate targets in the activin pathway or more effective therapies in PDAC cachexia.

## Results

### Activins are differentially expressed in pancreatic tumor and tumor-derived cell lines in murine PDAC cachexia models

Three cell lines, KPC32043, KPC32047, and KPC32908 derived from a genetically engineered PDAC mouse model LSL-**K**ras^G12D^;LSL-Tr**p**53^R172H^;Pdx1-Cre (hereafter referred to as KPC), were analyzed with RT-qPCR for expression of two activin/inhibin β subunit genes, *Inhba* and *Inhbb*. These cell lines expressed increased levels of *Inhba* mRNA, in 52- (p<0.001), 132-, and 551-fold (p<0.0001), respectively, compared to normal mouse pancreas (Figure 1A), while *Inhbb* is undetectable (data not shown). The three KPC cell lines express significantly different level of *Inhba* with highest level by KPC32908 (Figure 1A).

**Figure 1.**
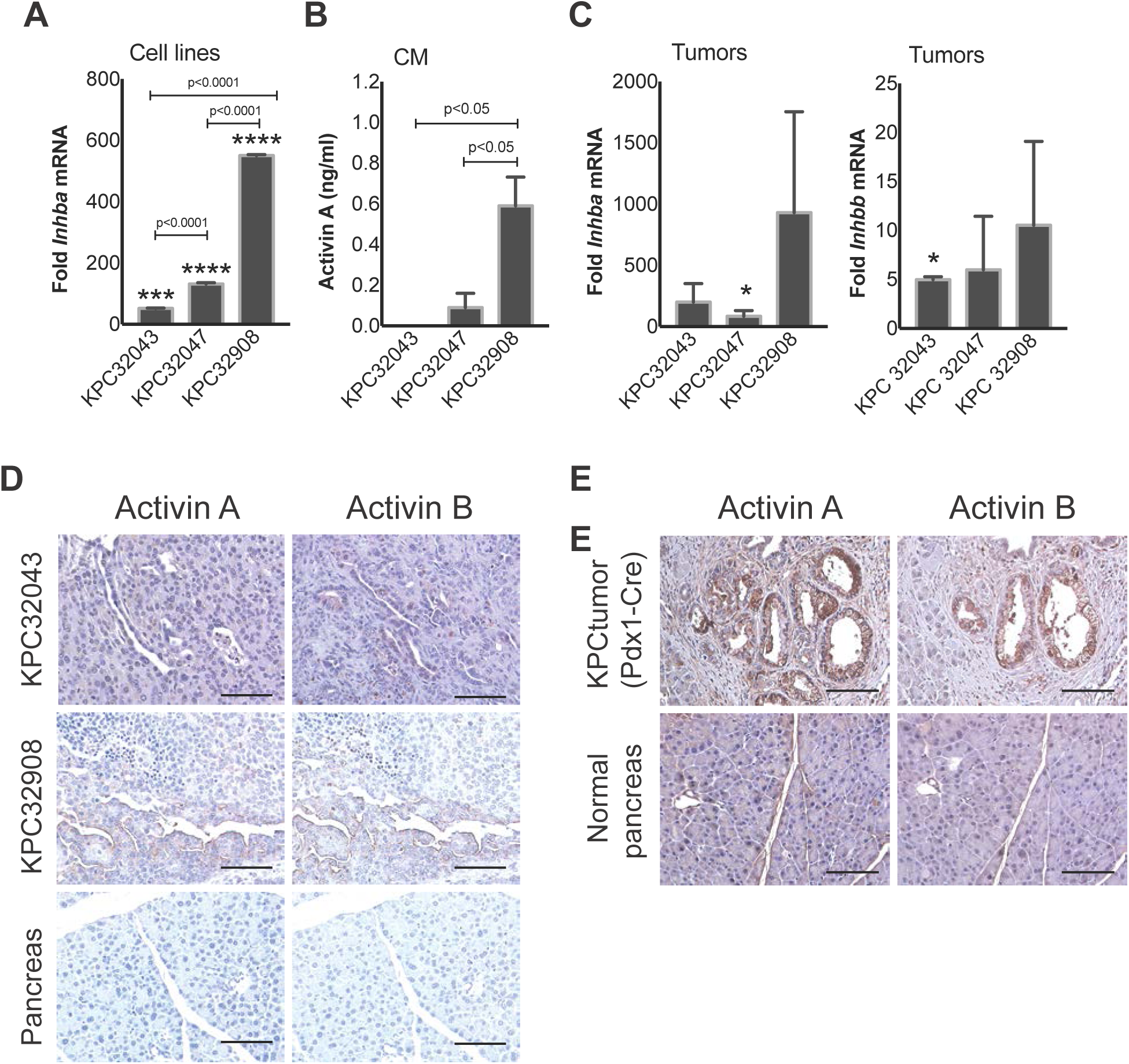
Differential expression of activin in pancreatic tumor and tumor-derived cell lines in murine and human pancreatic ductal adenocarcinoma (PDAC) cachexia models. A Quantitative RT-PCR demonstrating varying levels of *Inhba* mRNA in cell lines: KPC32043, KPC32047, and KPC32908 that were derived from the pancreatic tumor of a genetically engineered mouse model (GEMM) carrying triple mutant alleles, LSL-**K**rasG12D;LSL-**p**53R172H;Pdx1-Cre (referred to as KPC). Data are presented as the mean fold change relative to normal mouse pancreas (± SD). B ELISA demonstrating activin A protein levels in the conditioned medium (CM) collected from the culture of the three KPC cell lines. Data are presented as Mean ± SD. C Quantitative RT-PCR demonstrating levels of *Inhba* and *Inhbb* mRNA in pancreatic tumors generated by injection of the KPC cell lines into the pancreas of C57BL/6J mice (hereafter referred to as KPC orthotopic model). Data are compared to normal pancreas of sham-operated mice and presented as Mean ± SEM (n = 3). D Immunohistochemistry (IHC) of pancreatic tumor tissues from two KPC orthotopic models (KPC32043 and KPC32908) probing activin A or activin B level. E IHC of pancreatic tumor tissues from GEMM KPC with Pdx1-Cre. Normal pancreas tissues were from the sex- and age-matched mice with wild-type (WT) Kras and WT p53 alleles. Scale bar: 100 µm. *, **, ***, and ****: Data were compared to normal pancreas. *P < 0.05, ***P < 0.001, ****P < 0.0001.

To determine whether the increased *Inhba* mRNA expression leads to an increase in the secretion of the βA subunit-containing activins, we collected conditioned medium (CM) from KPC cell line cultures and measured the activin A level with the enzyme-linked immunosorbent assay ELISA method. The result showed that KPC cancer cells secreted activin A into the culture medium with the KPC 32908 CM containing more activin A than the KPC32047 by 6.6-fold (p<0.05) and the KPC 32043 CM (undetectable, p<0.05) (Figure 1B). The increased activin A could be due to the increase in the mRNA levels (Figure 1A). These results indicate that KPC32908 tumor likely releases more activin A and other factors into blood system than the KPC32043 tumor, which would have greater impact on the host organ biology.

We went on examining *Inhba* and *Inhbb* mRNA expression in an orthotopic model of PDAC cachexia that we developed by injecting the KPC cell lines into the pancreas of wild-type (WT) C57BL/6J mice. Compared to the normal pancreas from sham-operated controls, the KPC32043, KPC32047, and KPC32908 tumors expressed high levels of *Inhba*, with increases of 201-, 85-, and 931-fold respectively though only KPC32047 reached statistical significance (Figure 1C, left). Low levels of *Inhbb* were detected in these cell lines with highest level up to 10-fold; however, KPC32047 and KPC32908 did not reach the significance (Figure 1C, right). This differential expression pattern in KPC tumor is generally in agreement with that displayed in the KPC cell lines (Figure 1A) showing high level of *Inhba* and low level of *Inhbb*.

To confirm activin proteins produced or present in the PDAC tumor tissue, we performed IHC staining with activin subunit antibodies. While only the *Inhba* mRNA, but not the *Inhbb*, is highly increased in the KPC cancer cell lines and the cell line-derived orthotopic tumors (Figures 1A and 1C), both the βA and βB protein subunits were detected in the tumor cells and the stromal tissues from the two orthotopic models (Figure 1D) and the KPC **g**enetically **e**ngineered **m**ouse **m**odel (GEMM) (Figure 1E).

### Activins are expressed by human PDAC tumors

Since the preclinical murine models increased the activins in their tumors (Figure 1), we wanted to further explore whether human PDAC tumors would have the similar IHC staining. As shown in the upper panel of Figure 2A, activin βA and βB subunits were also detected in a panel of tumor sections from PDAC patients enrolled in our pancreatic cancer cachexia study (shown are representative images), compared to the normal pancreas placed at the end of the panel.

**Figure 2.**
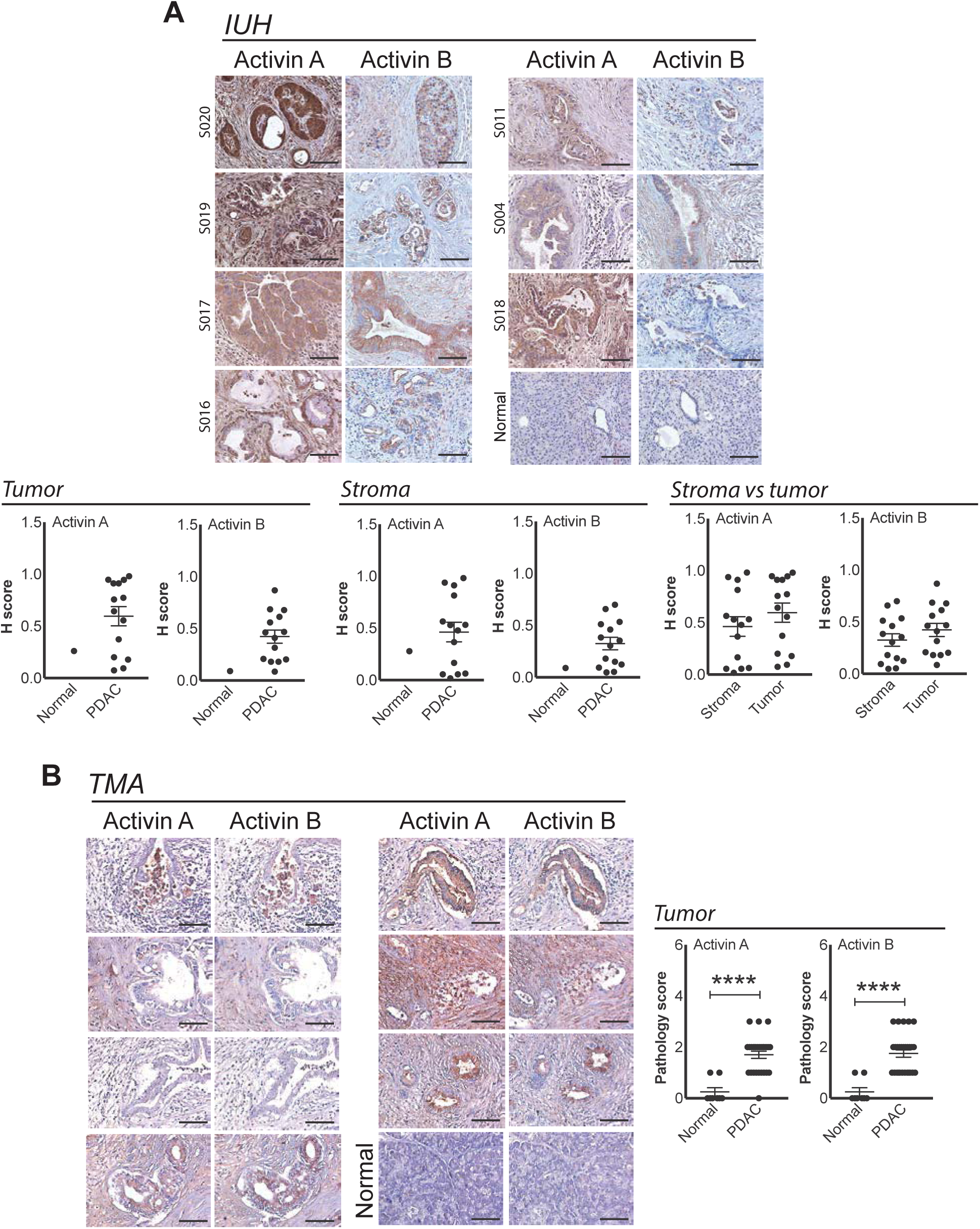
Activin proteins are expressed by human PDAC tumors. A IHC of tumor tissues from PDAC patients enrolled in the Indiana University Hospital (IUH). Shown are representative images. IHC quantification data from 16 PDAC patients are shown in the lower panel. B IHC of PDAC cancer tissue microarray (TMA) purchased from the US Biomax, Inc. IHC quantification data of normal pancreas (n=8) and PDAC tumor (n=27) are shown in the right panel. Data are presented as Mean ± SEM. Scale bar: 100 µm. ****P < 0.0001.

After the tissue sections were imaged with the Aperio whole slide digital imaging system at 20x, H-scores were generated to compare activin βA or βB subunit expression levels in normal pancreas versus tumor (Figure 2A, lower left panel). As only one normal pancreas tissue sample was available, the statistical analysis was difficult but the average levels of both βA and βB had a trend towards being higher in PDAC tumor or PDAC stroma (Figure 2A, lower middle panel) than in the normal pancreas counterparts. Furthermore, the average level of staining is not significantly different between tumor and stroma, though both show wide ranges of staining intensities from highly stained to lightly stained (Figure 2A, lower right panel). This variation may reflect the molecularly and pathologically heterogeneous nature of PDAC.

Because of the difficulty to obtain normal pancreas tissues, we purchased the Human Pancreatic Cancer Tissue Microarray (TMA) where more normal pancreas tissues are provided. These tissue spots were stained with the same antibodies and staining protocol as for the tissues from our own patient cohort but quantification was performed using a pathology hand score method by three pathologists, instead of using the H scoring system. Shown in Figure 2B are representative images on the left panel and the quantification graphs on the right panel. The staining result showed that activin expression is significantly increased in PDAC tumor compared to the normal pancreas (p<0.0001).

### *INHBA*, but not *INHBB* correlates with reduced survival of human PDAC patients identified by TCGA query

The data described above indicate that activin A may play an important role in PDAC cachexia. Indeed, query of the Oncomine database for studies in pancreatic cancer demonstrates 3 to 33-fold increases in *INHBA* gene expression versus controls in tumor (Figure 3A); *INHBB* showed no significant results. Furthermore, query of the Cancer Genome Atlas (TCGA) pancreatic cancer database for tumor expression of *INHBA* and related protein family members identified a correlation of *INHBA* with reduced overall survival of patients but not for *INHBB*, *MSTN*, *GDF11* and their inhibitors (Figure 3B).

**Figure 3.**
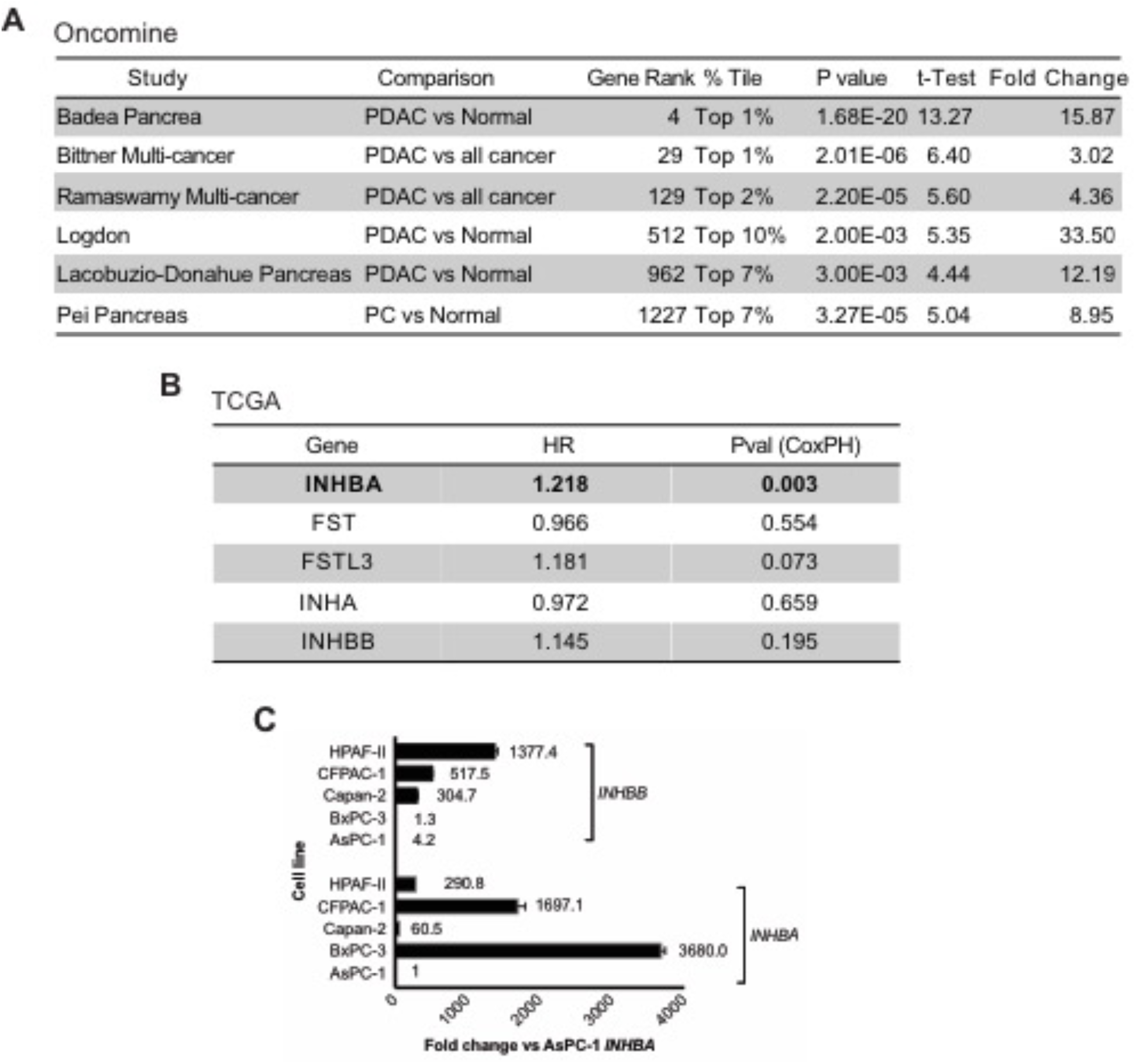
Elevated expression of *INHBA*, but not *INHBB* in Oncomine analysis of human PDAC tumors; *INHBA* expression, but not *INHBB* correlates with reduced survival in TCGA. A Query of the Oncomine database for studies in pancreatic cancer. B Query of the TCGA pancreatic cancer database for *INHBA* and related family members. C Quantitative RT-PCR probing *INHBA* and *INHBB* expression in human PDAC cell lines to support our database query results. The level of *INHBA* and *INHBB* expression are normalized to one (AsPC-1) of the five cell lines.

These database query results were supported by our further analysis of *INHBA* and *INHBB* expression in human PDAC-derived cell lines purchased from ATCC. For *INHBA* expression level, four of the 5 cell lines examined express the gene at an average level of 1146-fold (ranging from 60.5- to 3680.0-fold) higher than the cell line, AsPC-1 that is set at 1 as a calibrator. For *INHBB*, one cell line (BxPC-3) expresses at a similar level (1.3 fold) to the calibrator AsPC-1 and the other four express an average level of 441-fold higher than the calibrator (ranging from 4.2 to 1377.4-fold) (Figure 3C). Thus, these human pancreatic cancer cell lines also display a pattern of higher *INHBA* expression than *INHBB*, reinforcing the role of *INHBA* in PDAC and PDAC cachexia.

### There is a systemic functional decline due to PDAC

The observations that the tumor in the PDAC cachexia murine model produces a high level of activin A and the cultured KPC cancer cells releases activin A into the medium (CM) led us to predict that the tumor-derived activin and other factors would cause a systemically biological response in PDAC. To test this prediction, we assessed PDAC-induced cachexia/muscle wasting development in the KPC32043 orthotopic tumor-bearing mice both over time and at the endpoint to provide a thorough picture of the kinetics.

The mouse body weights were monitored over a period of 22 to 23 days after the mice were implanted with the KPC cells into their pancreas. We observed that the relative body weights started to drop at approximately 12 days after implantation and the drop continued till the mice reached the pre-specified point for euthanizing and organ collection. On the other hand, the sham-operated mice steadily gained weights up to a difference of approximately 9% body weight higher than KPC mice at the end of the experiment (both sham and KPC mice were normalized to their initial body weight (IBW) measured at the beginning of the experiment) (p<0.001) (Figure 4A).

**Figure 4.**
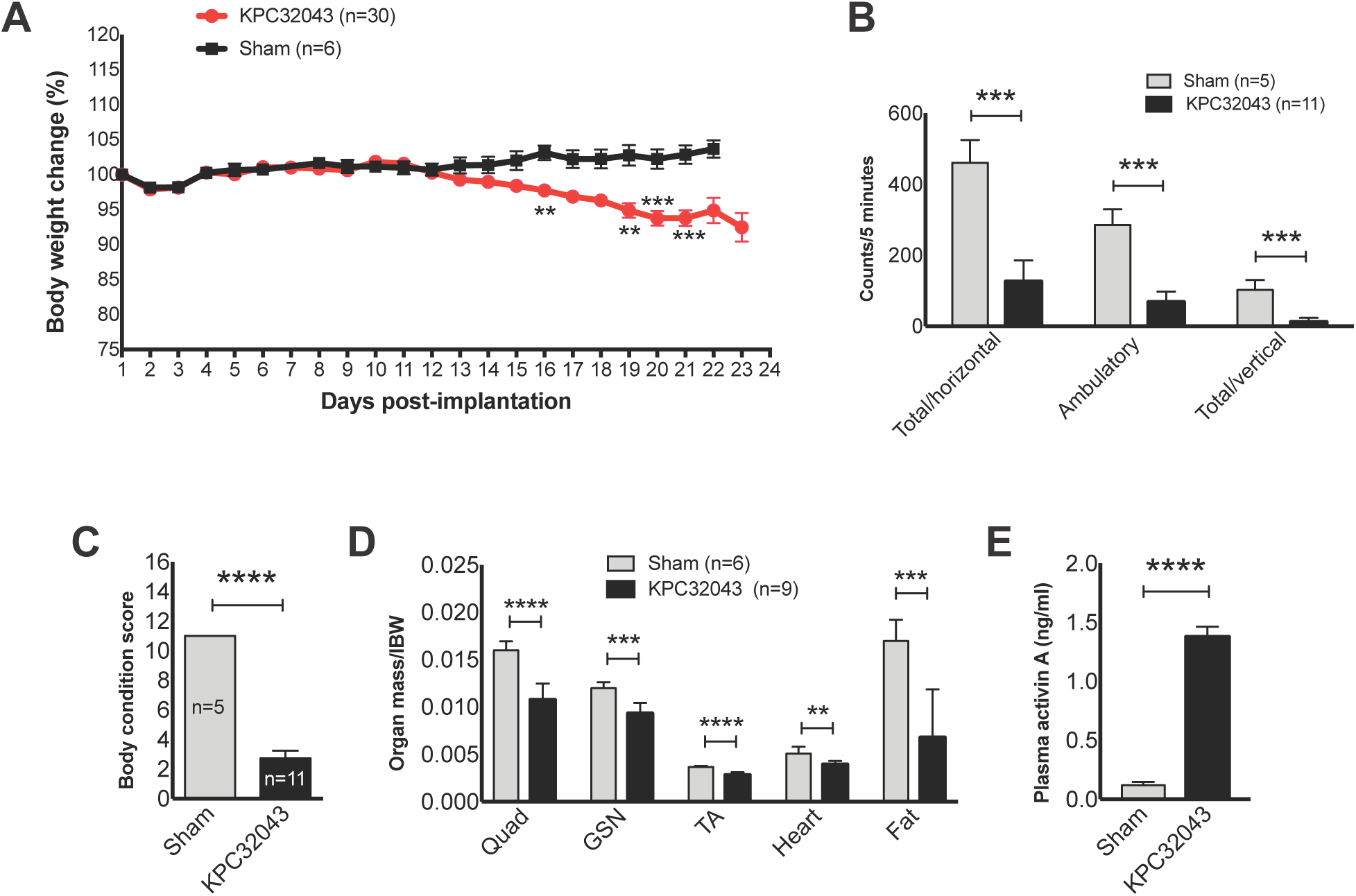
There is a systemic functional decline due to PDAC. A-D Systemic changes in body weight, activity, body condition, and organ weight in response to PDAC tumor developed by injection of KPC32043 cancer cell line, compared to the sham-operated mice. E Elevated activin A level in plasma from the tumor-bearing mice as determined with ELISA. Data are presented as Mean ± SEM. **P < 0.01, ***P < 0.001, ****P < 0.0001.

Before euthanizing, the mice were evaluated for activity and body condition. The tumor-bearing mice displayed significant reduced activity as determined by monitoring of the horizontal and vertical movements of mice within the acrylic chambers of VersaMax AccuScan activity monitors (Figure 4B). The body condition was markedly declined due to the tumor growth, compared to that of the sham controls (p<0.0001) (A score 11 is normal and 5 is consistent with imminent death) (Figure 4C). Weights of the tumor-bearing mouse organs harvested at the endpoint were significantly lower than the sham’s (Figure 4 D).

Since the KPC tumor-derived cell lines release activin A into the culture medium as shown in Figure 1B, the tumor would be expected to release activin A into the blood system and to be, at least partly, responsible for the cachectic syndrome described above. We measured activin A level in plasma from KPC32043 orthotopic mice and detected an 11.5-fold increase compared to the sham control (p<0.0001) (Figure 4E).

### There is a systemic activin response to PDAC

To assess the biological response of the host to the PDAC tumor at molecular level, we investigated activin mRNA expression in multiple organs. *Inhba* mRNA was significantly induced in most of the organs examined including spleen, heart, fat and kidney with a downregulation in liver and skeletal muscle, compared to the sham mice (Figure 5A). Notably, we found that *Inhbb* was also significantly induced in all the organs (Figure 5B). Generally, activin subunits are increased in the organs analyzed (Figure 5C).

**Figure 5.**
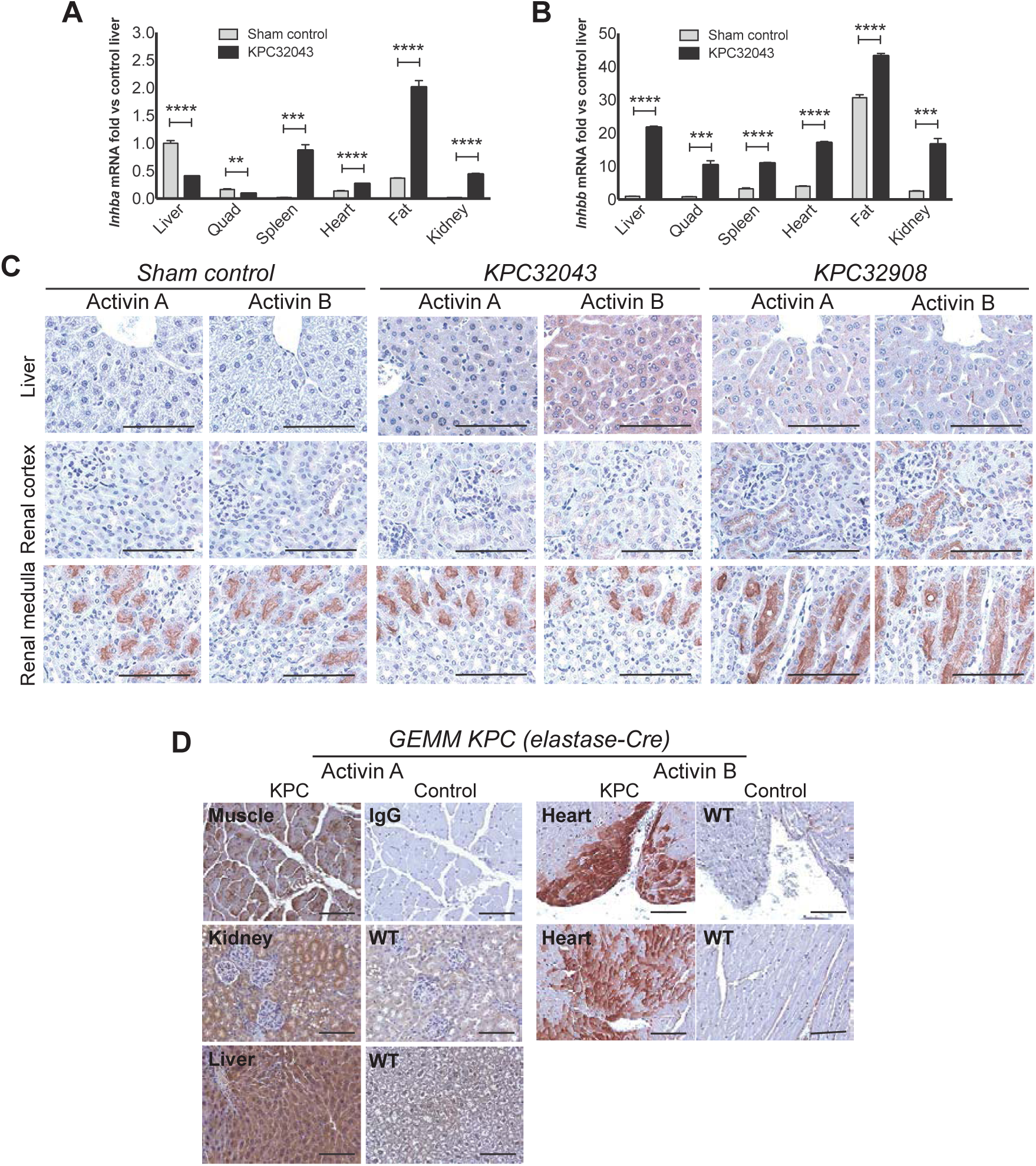
There is a systemic activin response to PDAC in murine KPC models. A-B *Inhba* or *Inhbb* mRNA expression in organs harvested at the endpoint from sham control and orthotopic KPC32043 tumor-bearing mice, determined by quantitative RT-PCR. C IHC of organs from two orthotopic KPC cancer cell injection models probing activin A or activin B expression. D IHC of organs from GEMM KPC (elastase-Cre) tumor-bearing mice probing activin A or activin B expression. Data are presented as Mean ± SEM. Scale bar: 100 µm.

We then further assessed activins in the KPC GEMM. As shown in Figure 5D, activin A (left panel) and activin B (right panel) are induced in multiple organs including muscle, kidney, liver or heart.

### PDAC tumor cells express factors that induce activins in and cause atrophy of muscle cells

Since muscle wasting is the feature of cancer cachexia and activins are elevated in muscle of the PDAC tumor mice, we asked whether the induction is a direct response to factors made by the PDAC tumor. To address this question, we exposed the murine muscle-derived C2C12 myoblasts and 4-day old differentiated myotubes to 75% of KPC32908 CM for 24 or 48 hours (schematic drawing in Figure 6A, upper panel) and analyzed *Inhba* and *Inhbb* mRNA expression with RT-qPCR. After CM treatment, both the *Inhba and Inhbb* were significantly increased in myoblasts and myotubes compared to their respective growth medium (GM)-treated controls (Figure 6A, lower panel). Myotube atrophy was marked induced after 48-h CM treatment (Figure 6B). These endogenous activins induced directly in response to PDAC tumor-derived factors would also contribute to the muscle wasting. To evaluate the degree of myotube atrophy, we measured the diameters of myotubes treated with 25%, 50% or 75% of CM for 48 hours. The average reductions in diameter were statistically significant at all dilutions in a CM dilution ratio-dependent manner; the higher the CM concentration, the more severe the atrophy (Figure 6B, lower panel).

**Figure 6.**
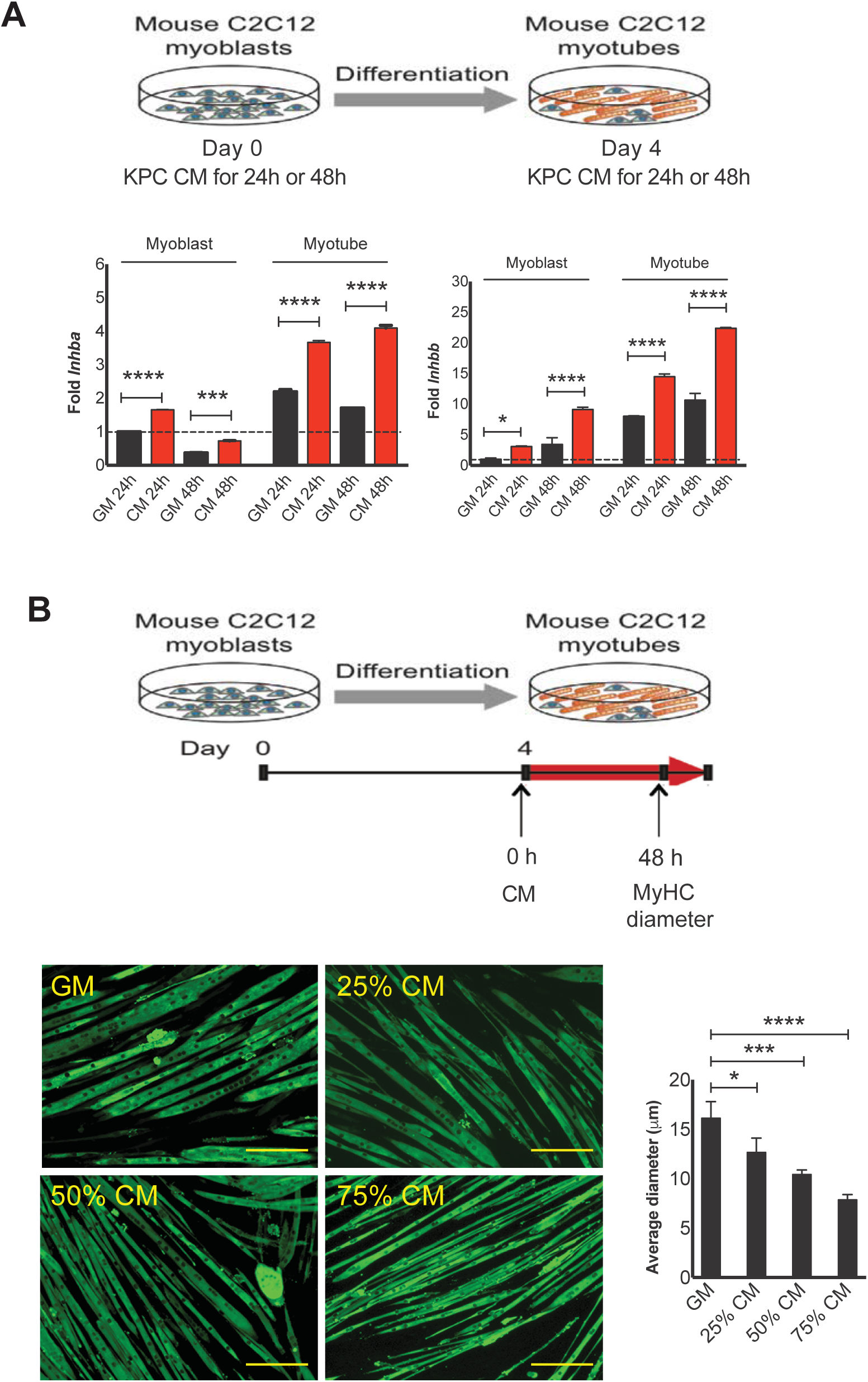
PDAC tumor cells express factors that induce activin expression in and cause atrophy of muscle cells. A Schematic drawing illustrating the approach of preparing myoblasts/myotubes and CM treatment (upper panel) and quantitative RT-PCR demonstrating the induction of *Inhba* and *Inhbb* by 75% of KPC32908 CM (lower panel). B Schematic drawing illustrating the approach of preparing myotubes and CM treatment (upper panel), images showing immunofluorescence staining of myosin heavy chain in myotubes to visualize myotube shape (lower panel, left), and graph showing average sizes measured from more than 200 of qualified myotubes per treatment condition (growth medium GM, 25%, 50%, or 75% of KPC32908 CM) (lower panel, right). *P < 0.05, ***P < 0.001, ****P < 0.0001. Scale bar: 200 µm.

### Activin inhibition in a murine model of KPC32043 PDAC cachexia reduces weight loss and prolongs survival

To determine whether blocking circulation activins would alleviate cachexia, we treated mice with ACVR2B/Fc. Intraperitoneal injection (i.p.) of ACVR2B/Fc at 10 mg/kg body weight on days 3, 10, 14 and 18 after surgery of KPC cell implantation reduced body weight loss with the effect starting at approximately two weeks after the first injection (Figure 7A, left). At the end of the experiment, the weight loss was completely prevented. ACVR2B/Fc treatment also improved body condition (Figure 7A, right), resulted in a retention of the hanging grid exploration activity (Figure 7B, left) and an increase in open field activity (Figure 7B, right), prolonged animal survival from median survival 25 days without treatment to 29.5 days with treatment (p<0.05) (Figure 7C), and reduction of the loss in skeletal and heart muscle as well as fat mass, though the effect on fat did not reach a significant difference (Figure 7D, left). Gene expression in muscle with the soluble receptor treatment did reduce *Atrogin-1* and *Murf-1* (Figure 7D, middle) as well as *Mstn*, but not the expression of *Inhba* or *Inhbb* (Figure 7D, right), suggesting this is not an activin endocrine loop.

**Figure 7.**
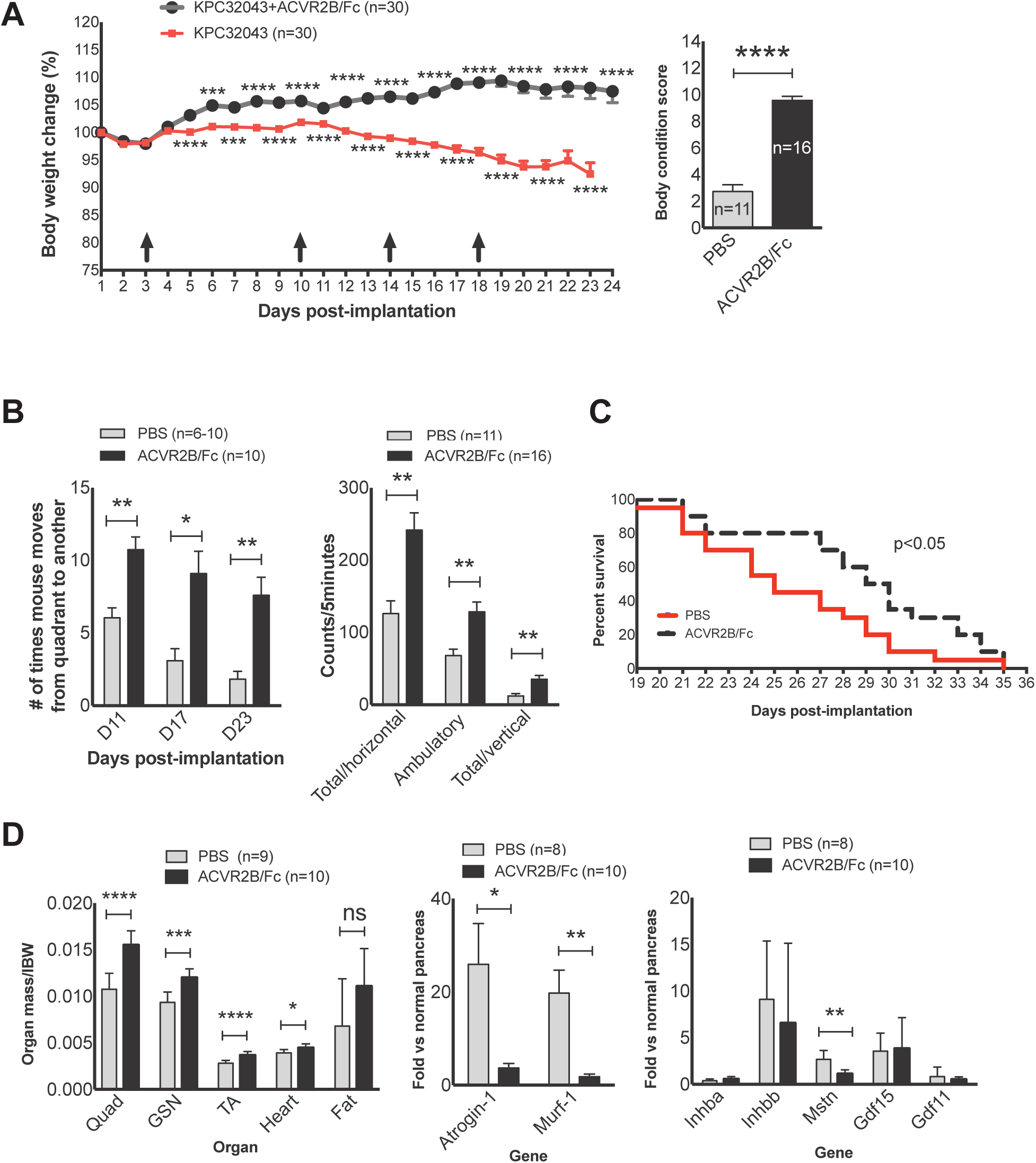
Activin blockade using ACVR2B/Fc reduces weight loss and prolongs survival in the orthotopic KPC tumor-cachexia mouse model. A Body weights recorded over time following KPC32043 cancer cell injection into the pancreas and ACVR2B/Fc administration (10 mg/kg body weight via i.p.) (left panel) and body condition scores at the end of the experiment (right panel). Arrow: Time points of ACVR2B/Fc administration. B Mouse activities measured at the indicated time points (left panel) or at the end of experiment (right panel) showing the therapeutic effect due to activin blockade. C Protective role of ACVR2B/Fc in mouse survival. D Effect of activin blockade on organ mass (left) and on the expression of protein degradation-related genes (middle) or activins and the related genes (right). Data are presented as Mean ± SEM. Arrow: Time points of ACVR2B/Fc administration. IBW: Initial body weight measured on the day of cancer cell injection. Data are presented as Mean ± SEM. *P < 0.05, **P < 0.01, ***P < 0.001, ****P < 0.0001.

### Activin inhibition with ACVR2B/Fc in a more aggressive, higher *Inhba*-expressing KPC32908 orthotopic mouse model is less effective

To explore whether the soluble receptor can exert a therapeutic role in a more aggressive, higher *Inhba*-expressing KPC32908 orthotopic mouse model, we i.p. injected the same does of ACVR2B/Fc as with the KPC32043 model on days 2, 7, and 12 after KPC32908 cell implantation and monitored the body weight. As with the KPC32043 model, ACVR2B/Fc treatment prevented body weight loss (Figure 8A); however, it did not prolong animal survival (Figure 8B).

**Figure 8.**
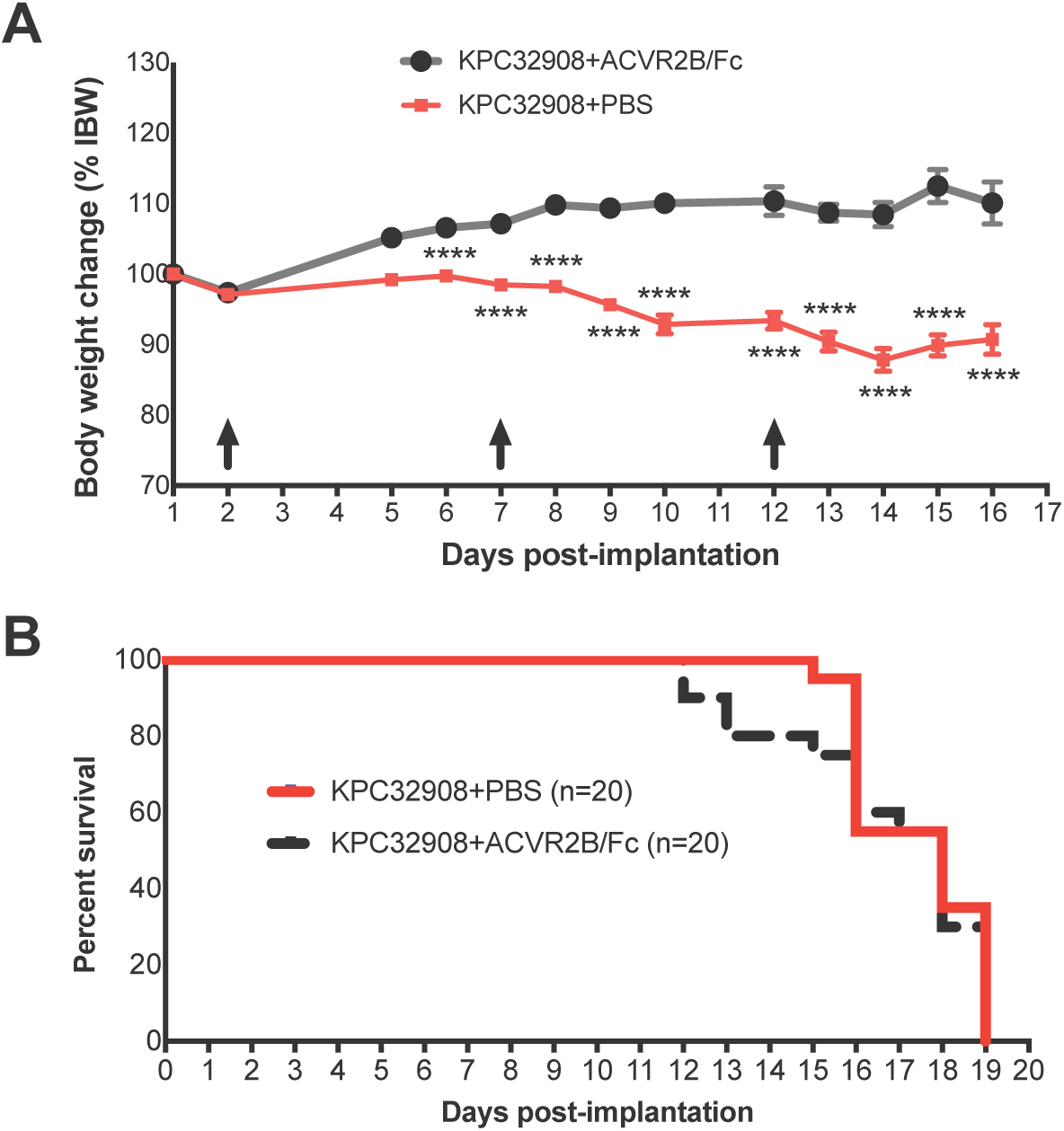
Body weight sparing but not prolonged survival with activin inhibition using ACVR2B/Fc in the more aggressive, higher *Inhba*-expressing orthotopic KPC32908 cachexia model. A Body weights recorded over time following KPC32908 cancer cell injection into the pancreas and ACVR2B/Fc administration (10 mg/kg body weight via i.p.). Data are presented as Mean ± SEM. Arrow: Time points of ACVR2B/Fc administration. B Survival of the KPC32908 tumor-bearing mice with or without ACVR2B/Fc administration. ****P < 0.0001.

### Skeletal muscle-specific dnACVR2B expression in MLC-dnACVR2B mouse model prevents PDAC-induced weight loss but does not improve survival

Since systemic administration of ACVR2B/Fc exerts an inhibitory effect on cachexia in the KPC orthotopic mouse model, we wanted to know how the ACVR2B-transduced activin signal in skeletal muscle contributes to PDAC muscle wasting. We implanted the KPC32043 cells into the pancreas of wild-type (WT) male C57BL/6 mice (WT+KPC32043) or MLC-dnACVR2B mice expressing skeletal muscle specific, dominant negative ACVR2B that lacks the kinase domain and thus blocks ACVR2B signaling (Tg+KPC32043). The weights of the mice were monitored daily. We observed that the PDAC tumor formation in mice with WT background induced steady weight loss over the 23 days after implantation (Figure 9A, left panel) and the difference of final body weight at sacrifice between the tumor-bearing mice and the sham controls reached 8.1% (p<0.01) (Figure 9A, right panel). However, ACVR2B signaling interruption in the MLC-dnACVR2B mice prevented body weight loss induced by KPC32043 tumor (Tg+ KPC32043), compared to the WT+KPC32043 (p<0.05) and the Tg sham, while there were no statistically significant differences in the body weights of WT sham versus transgene (Tg) sham over the whole period of experiment (Figure 9A). These observations suggest that dnACVR2B expression does not affect mice under physiological condition but the ACVR2B-mediated signaling is indeed important for the development of muscle wasting under a pathological condition like PDAC.

**Figure 9.**
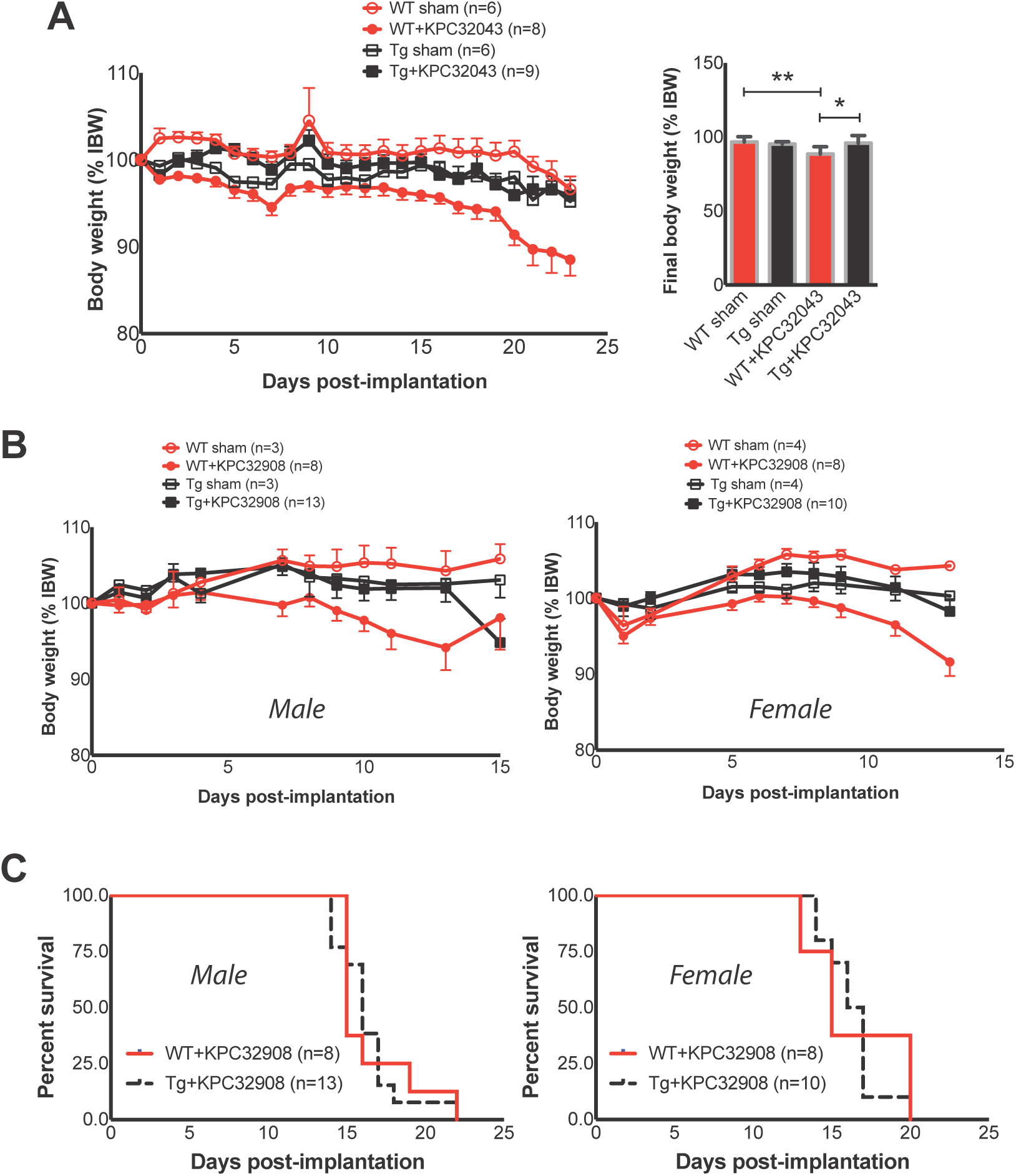
Skeletal muscle-specific dnACVR2B expression in MLC-dnACVR2B mouse model prevents PDAC-induced weight loss but does not improve survival. A Body weights of KPC32043 pancreatic tumor-bearing wild-type (WT) C57BL/6 mice or Tg (transgenic representing MLC-dnACVR2B) mice, normalize to each mouse’s own IBW (left); final body weight (right). B Body weights of KPC32908 pancreatic tumor-bearing WT C57BL/6 mice or MLC-dnACVR2B mice, normalize to each mouse’s own IBW. C Survival of the KPC32908 pancreatic tumor-bearing mice with WT background or MLC-dnACVR2B, monitored from the day of KPC cell injection to the day when the mice reach a moribund endpoint. IBW: Initial body weight measured on the day of cancer cell injection. Data are presented as Mean ± SEM. *P < 0.05, **P < 0.01.

We further evaluated whether interruption of ACVR2B signaling would prevent body weight in the transgenic mice bearing the more aggressive KPC32908 tumor. As shown in Figure 9B, body weight loss was prevented in Tg+KPC32908 mice compared to the WT+KPC32908 and the Tg sham. However, survival was not prolonged by dnACVR2B expression (Figure 9C) (p>0.05). Similar results were seen in males and females. Overall these results suggest that activin signaling in other tissues outside muscle must play a role in cachexia.

## DISCUSSION

Activin A is best known for its function in embryogenesis and reproduction with increasingly demonstrated roles in tumorigenesis. However, its role in cancer cachexia has just begun to be appreciated. In the present study, we have identified a systemic organ activin response to PDAC tumor in murine and human models of PDAC cachexia. While the tumor expressed a high level of activin A, multiple organs expressed both activin A and activin B in response to the tumor-derived factors. Furthermore, we have demonstrated that the tumor-derived factors are causally involved in muscle wasting as direct exposure of muscle-derived C2C12 myotubes to PDAC cancer cell-derived conditioned medium significantly reduced myotube size, an experimental model mimicking the *in vivo* muscle wasting. Additionally, blocking activin with the soluble receptor ACVR2B/Fc reduced PDAC cachexia but prolonged survival only in a less-aggressive mouse model. Finally, skeletal muscle-specific dnACVR2B expression in the MLC-dnACVR2B mouse model prevented PDAC-induced weight loss but did not improve survival. These results suggest that a systemic activin-targeting therapeutic strategy will be necessary in combination with anti-cancer agents and further developing new targeted therapeutics based on the sophisticated mechanisms underlying PDAC cachexia will be urgently needed to effectively combat the wasting syndrome in PDAC patients.

Our observation that the PDAC tumor-released factors induced expression of activins including activin A and activin B in multiple host organs warrants targeting both the circulating activin and the induced endogenous activin to achieve stronger inhibition on the activin-mediated cachectic signaling. Although the soluble activin receptor ACVR2B/Fc can block the circulating activins, it cannot prevent the endogenous activin from accessing the receptor and initiating the signaling due to its lack of the ability to bind to the cell surface heparin-sulfated proteoglycans (Nakamura et al, 1991; Sidis et al, 2006). The induced endogenous activins may indeed play an important part in muscle wasting through an autocrine manner. A recent published work supports this concept, which showed that mouse leg muscle mass was significantly increased due to the inhibition of endogenous activins by ectopic expression of engineered inhibitory activin A or activin B-carrying rAAV6 vectors (Chen et al, 2015).

It has been shown that activin A can induce the expression of *INHBB* subunit mRNA (Eramaa et al, 1995). One could argue that the organ activins induced in response to PDAC tumor were the result of exogenous activin stimulation. We observed that while ACVR2B/Fc administration alleviated cachexia in the orthotopic mouse model, it did not change the expression levels of *Inhba* and *Inhbb* in the organs examined. This suggests that the organ activin induction is not an activin endocrine loop; instead, other factors released by the tumor are responsible for the induction. Identification of these factors would offer the means to prevent the endogenous activin induction.

ACVR2B/Fc has broader spectrum of ligand binding including myostatin, activin, GDF11 and other TGF-β family ligands, which has raised a concern of off-target effects. To overcome this issue, Chen et al developed antagonists specific to activin A or activin A and B (Chen et al, 2015; Makanji et al, 2011). They found that the use of the antagonist toward both activin A and activin B resulted in a greater increase in muscle mass than the one toward activin A only. Therefore, our finding of systemic induction of activin B, in addition to activin A, in PDAC cachexia is significant. In fact, while activin A was induced in some of the organs we examined, activin B was consistently induced in all (Figure 5 A and B). Thus, we suggest that endogenous activin B-targeting agents should be considered in a PDAC cachexia therapeutic strategy. It should be mentioned that, like the ACVR2B/Fc, the antagonists that Chen et al generated also only block the circulating activins but not the endogenous ones (Chen et al, 2015). Therefore, new generations of antagonists need to be designed to be effective on the endogenous activins.

In the present study, we treated two models of KPC tumor-bearing mice with ACVR2B/Fc, one with lower *Inhba*-expressing (KPC32043) tumor and the other with higher *Inhba*-expressing (KPC32908) tumor. ACVR2B/Fc was less effective in the KPC32908 mice than the KPC32043 mice in terms of survival benefit. This result suggests that KPC32908 tumor likely releases more activin A, which may simply require higher dose of ACVR2B/Fc to be effective. Alternatively, higher level of *Inhba* expression in tumor may confer a more aggressive feature on tumor itself and this would in turn post more profound impact on the host, such as inducing more endogenous activins in organs; while ACVR2B/Fc cannot antagonize the endogenous activins, as reasoned above (Chen et al, 2015; Nakamura et al, 1991; Sidis et al, 2006).

Muscle tissue represents more than 40% of body weight and thus its wasting accounts for a larger part of cachectic body weight loss. Despite this, other tissues/organs such as fat and heart are recently suggested to play an important role in the cachexia development and may be responsible for muscle wasting (Argiles et al, 2015). In our PDAC models, mass loss of multiple organs was evident (Figure 4D). Furthermore, activin induction was observed in multiple organs (Figure 5 A and B). Our data obtained by using the MLC-dnACVR2B mice expressing skeletal muscle specific, dominant negative ACVR2B further support the roles played by other organs, which showed that the weight loss was prevented in the MLC-dnACVR2B mice bearing the orthotopic KPC tumor compared to the WT mice with the tumor (Figure 9 A and B); however, the dnACVR2B expression in skeletal muscle did not show survival advantage (Figure 9C), indicating that activin signaling in other tissues outside muscle contributes to the cachexia.

In summary, the present study demonstrates PDAC tumor induces cachexia and functional decline through producing and releasing activins and other cachectic factors into the circulation that causes a systemic biological response from multiple distant organs of mouse as well as human patients. The systemically induced endogenous/autocrine activins, together with the exogenous/endocrine (circulating) activins might both contribute to the development of muscle wasting and cachexia. The soluble activin receptor ACVR2B/Fc has a limited role in blocking the induced organ activins and the tumor-secreted, activin-inducing factors have yet to be identified. These observations and postulations are illustrated in Figure 10. Finally, developing new agents that can block both sources of activins will be needed to provide more effective therapy for PDAC cachexia.

**Figure 10.**
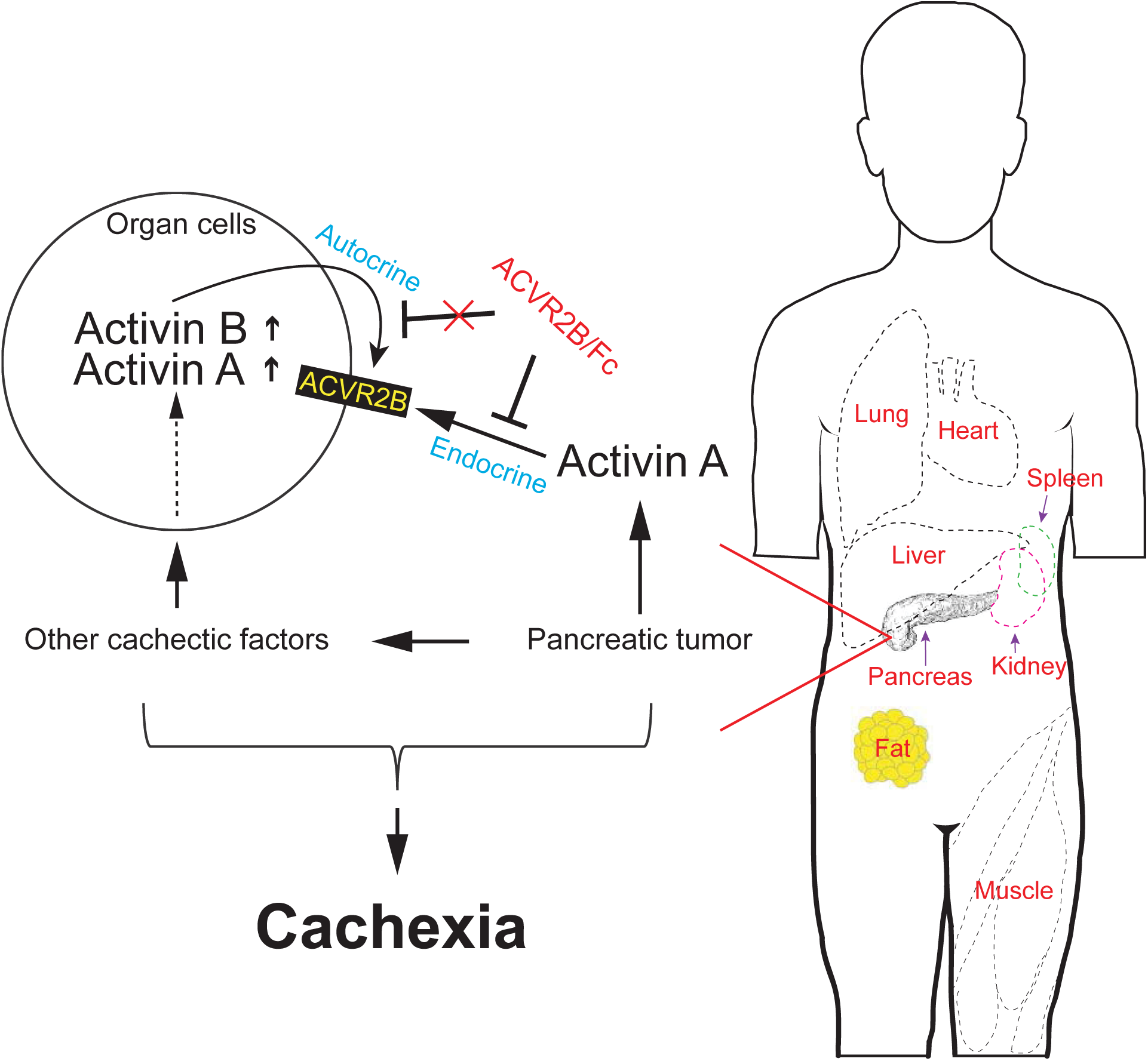
Proposed framework illustrating the systemic activin induction in response to pancreatic cancer. PDAC tumor releases activins and other cachectic factors into the circulation, which induces a systemic biological response from multiple distant organs of mouse as well as human patients. The tumor-derived and host-derived activins and other factors together result in PDAC cachexia. The soluble activin receptor ACVR2B/Fc can block the endocrine activins but has a limited role in autocrine activins, which warrants effective therapeutics targeting both endocrine and autocrine actions of activins.

## Materials and Methods

### Human and murine cell lines

Human pancreatic cancer cells, AsPC-1, BxPC-3, Capan-2, CFPAC-1 and HPAF-II (ATCC, Manassas, VA, USA) were maintained in the respective media including RPMI1640, McCoy’s 5a Medium, Iscove’s Modified Dulbecco’s Medium, or Eagle’s Minimum Essential Medium as specified by ATCC. Genetically engineered mouse-derived pancreatic cancer cells, KPC32043, KPC32047, and KPC32908 (kind gifts from Dr. David Tuveson at Cold Spring Harbor Laboratories) and murine skeletal muscle-derived C2C12 myoblasts (ATCC) were maintained in DMEM medium. All media were supplemented with 10% fetal bovine serum and 1%penicillin/streptomycin.

Myoblast differentiation to myotubes was induced by shifting confluent cultures to DMEM supplemented with 2% horse serum (named differentiation medium, or DM) and the DM was replaced every day for 6 days.

### Animal experiments

All animal studies were approved by the Institutional Animal Care and Use Committee at Indiana University School of Medicine and in compliance with the National Institutes of Health Guidelines for Use and Care of Laboratory Animals.

Cancer cachexia murine orthotopic model: The murine KPC pancreatic cancer cells (5 × 10^5^/mouse) were injected orthotopically into the pancreas of ~10-week old wild-type (WT) C57BL/6 mice purchased from Jackson Laboratory.

Genetically engineered mouse (GEM) model (GEMM): The GEMM **K**-rasG12D; **p**53R172H;Pdx1-**C**re (KPC) was generated by crossing LSL-K-rasG12D mouse with LSL-p53R172H mouse (obtained from the mouse repository of National Cancer Institute; NCI), producing LSL-K-rasG12D;LSL-p53R172H (i.e., KP), which was then crossed with Pdx1-Cre (NCI) that expresses a pancreas-specific Cre recombinase, giving rise to KPC that would develop tumor at the pancreas. The KPC model mimics human PDAC by introducing a high degree of genomic instability due to point mutations in the oncogene Kras and the tumor suppressor gene p53 (Hingorani et al, 2005).

The pancreatic tumor and organ tissues of GEMM KPC mice with elastase promoter-driven Cre expression in pancreas were provided by Stephen Konieczny at Purdue University.

Murine MLC-dnACVR2B model: The mice (a gift from Se-Jin Lee at Johns Hopkins University), which express skeletal muscle-specific, dominant negative ACVR2B lacking the kinase domain, were implanted with the murine KPC pancreatic cancer cells (5 × 10^5^/mouse) at age of 10 weeks.

Mice were weighed daily on the following day of injection and thereafter for approximately 2-3 weeks and euthanized under isoflurane anesthesia when they reached the pre-designated endpoint. Body composition, mouse activities, and muscle strength were monitored weekly or at the end of experiments (see below for details).

The endpoint determination for euthanizing was dependent on the aims of studies: For study of therapeutic intervention effects on survival in cancer, mice were allowed to become moribund; and for study of host biological response to tumor/therapeutic intervention, mice were allowed to reach “tissue endpoint”. We used a body posture, activity level and eye appearance-based scoring system plus an additional supporting criterion (see later) to determine the “moribund” or “tissue endpoint”: Mice were euthanized at a score of 8 or 9 for moribund and 7 or 8 for tissue endpoint (The higher the score, the sicker the mouse, with the highest score being 9). The supporting criterion is that tumor-bearing mice would be euthanized if they fail to move or resist after gentle manipulation.

Body composition measurement: A Quantitative Magnetic Resonance (QMR) technology-based EchoMRI instrument (EchoMRI, LLC, Houston, TX) was used to track changes in body composition (fat mass and lean mass) over a period of time in live, non-anesthetized, non-sedated mice. The mouse was placed into a holding tube and the tube was inserted into the instrument, followed by scanning. After scanning, the tube containing the mouse was removed from the instrument and the mouse was returned to the home cage. The duration of a single EchoMRI measurement is less than one minute.

Mouse activity measurement: Activity was assessed using non-invasive monitoring of horizontal and vertical movements by placing mice within the acrylic chambers of VersaMax AccuScan activity monitors and recording for 5-30 min.

Grip strength: This test measures maximal muscle strength of limbs. The mouse was allowed to grasp a wire mesh grid with its paws and then was gently pulled backward by its tail until it released the grid. This procedure was repeated three times with at least several-minute interval between two tests. Measurements were recorded in grams by a force transducer attached to the grid.

Hanging time test: The test measures combined forepaw and hindpaw strength. The mouse was placed on a wire grid which was then inverted over a foam pad. The duration of hanging to fall was recorded.

Activin inhibition: We utilized ACVR2B/Fc, a soluble form of the activin type IIB receptor fused to an Fc domain to block the binding of activin and related ligands to the common ACVR2B receptor. C57BL/6 male mice bearing orthotopic KPC tumors were treated with either PBS (vehicle control) or ACVR2B/Fc (10mg/kg) at the pre-designated intervals.

Tissue collection: At the end of experiments, mice were euthanized and tissues were collected and weighed, followed by snap frozen in liquid nitrogen, freezing in cold 2-methylbutane, or fixation in 10% neutral buffered formalin solution, and then placed in −80^°^C for longer term storage.

### Quantitative RT-PCR

Total RNA was extracted using the QIAzol Lysis Reagent and miRNeasy Mini Kit (Qiagen) and reverse-transcribed into complementary DNA (cDNA) using the TaqMan Reverse Transcriptase Reagents (Life Techlologies) or the Verso cDNA Synthesis Kit (Thermo Scientific), following the manufacturer’s instructions. Equal amounts of cDNA were subjected to Quantitative real-time PCR performed using the TqaMan Universal Master Mix II with UNG on the LightCycler Instrument (Roche). All the gene-specific TaqMan PCR-based Gene Expression Assays were purchased from Life Technologies. Obtained CT values for target and housekeeping genes were used to calculate relative transcript abundance using the ΔΔCT method. Data were presented as the mean fold change.

### Immunohistochemistry (IHC)

IHC was performed on 5 μm formalin-fixed, paraffin embedded (FFPE) tissue sections. The sections were first deparaffinized and rehydrated, followed by heating in 0.01 M citrate buffer (pH 6.0) to retrieve antigens. Endogenous peroxidase was eliminated by incubation in 3% H2O2. The sections were incubated with the specific primary antibodies to INHBA from R&D Systems or to INHBB from antibodies-online at 1:100 and then with peroxidase conjugated secondary antibodies, ImmPRESS anti-goat or anti-rabbit from Vector Laboratories. The immunostaining was visualized with ImmPACT DAB from Vector Laboratories. Thereafter, the sections were conterstained with Hematoxylin.

### IHC stain imaging and quantification

Images were acquired at 20x using a color (MRc) Zeiss AxioCam camera mounted on a Zeiss Axio Observer.Z1 inverted microscope.

For image quantification, whole slide images were taken using the Aperio whole slide digital imaging system. The Aperio Scan Scope CS system was used (360 Park Center Drive, Vista, CA 92081). The system imaged all slides at 20x. The scan time ranged from 1 ½ minutes to a maximum time of 2.25 minutes. The whole images were housed and stored in their Spectrum software system and images were shot from the whole slides.

The Positive Pixel Count algorithm was used to quantify the amount of a specific stain present in a scanned slide image. A range of color (range of hues and saturation) and three intensity ranges (weak, positive, and strong) were masked and evaluated. The algorithm counted the number and intensity-sum in each intensity range, along with three additional quantities: average intensity, ratio of strong/total number, and average intensity of weak positive pixels. The algorithm had a set of default input parameters when first selected—these inputs have been pre-configured for Brown color quantification in the three intensity ranges (220-175, 175-100, and 100-0). Pixels, which were stained but did not fall into the positive-color specification, were considered negative stained pixels—these pixels were counted as well, so that the fraction of positive to total stained pixels was determined.

### Conditioned medium (CM) treatment

CM was collected from confluent culture of KPC pancreatic cancer cells and added to C2C12 myoblast or 4-day old differentiated myoytube culture at 25%, 50%, or 75% concentration in DM. The culture containing the CM was continued for additional 24 or 48 hours and RNA was extracted from the cells using the standard method as described above. Myotubes were also used for analysis with immunofluorescence microscopy.

### Immunofluorescence microscopy

C2C12 myotubes cultured in 12-well plates were fixed in 4% paraformaldehyde and permeabilized in 1% Triton X-100 at room temperature, followed by blocking in Sea Block Blocking Buffer (Thermo Fisher Scientific) supplemented with 0.2% Tween 20. The myotubes were then incubated with the myosin heavy chain (MyHC) primary antibody (DSHB) at 1:100 dilution and subsequently with the Alexa Fluor 488-conjugated anti-mouse secondary antibody (1:1000 dilution; Invitrogen). Nuclei were stained with 1 µg/ml 4′,6-diamidino-2-phenylinole (DAPI) (Calbiochem). Images were acquired at 10x using a monochrome (MRm) Zeiss AxioCam camera mounted on a Zeiss Axio Observer.Z1 inverted microscope.

### Myotube size quantification

The Alexa Fluor 488-labeled myotubes were assessed for the size changes following CM treatment. Fifteen fields from each well were randomly selected at imaging acquisition and three wells per experimental condition gave rise to 45 fields from which a total of approximately more than 200 qualified myotubes were measured. The average diameter of the myotubes per condition was calculated and expressed as Mean ± SD.

### ELISA

Activin A ELISA was performed using the Activin A ELISA kit (R&D Systems), per manufacturer’s protocol. In brief, 96-well plates were coated with the capture anti-activin A antibody overnight at room temperature and nonspecific binding was blocked with 1xdiluent followed by sample (conditioned medium or mouse plasma) addition and incubation. Biotinated anti-activin A detection antibody was added, which were subsequently incubated with Streptavidin-HRP. The Substrate Solution was added to produce a visible signal and the optical density of the signal was determined using a microplate reader (BioTek^®^ Instruments, Inc.).

### ACVR2B/Fc preparation

The ACVR2B/Fc fusion protein was stably expressed in the Chinese hamster ovary cells (a gift from Se-Jin Lee at Johns Hopkins University), purified from the conditioned medium using a protein A Sepharose (GE Healthcare), and dialyzed against 1 x phosphate-buffered saline (PBS) using Slidealyzer dialysis kit. The dialyed protein was confirmed by Western Blot along with the recombinant ACVR2B/Fc protein from R&D Systems as a positive control.

### Statistical analysis

Comparisons between values were performed using a Student’s t-test. Comparison of multiple groups, an ANOVA coupled with Tukey’s post-hoc test was applied. For all statistical analyses, the level of significance was set at a probability of <0.05.

## Acknowledgements

This work was funded in part by grants to TAZ from NIH (grants R01CA122596, R01CA194593, and R01GM092758), and to LGK from NIH (R01DK096167), the Lilly Endowment, Inc., the IU Simon Cancer Center, the Lustgarten Foundation, and the IUPUI Signature Center for Pancreatic Cancer Research.

## Author contributions

Study conception and design: XZ, MP, YL, LGK, TAZ

Acquisition of data: XZ, MP, CP, YJ, JL, GES, GJ, TAZ

Drafting of manuscript: XZ

Critical revision: XZ, MC, LGK, TAZ

